# Does non-Mendelian chromosome transmission and unusual sex determination affect male mate choice in the fly *Bradysia coprophila*?

**DOI:** 10.1101/2024.12.30.630734

**Authors:** Christina N Hodson, Robert Baird, Maddy Hodgemen, Shona Dury, Laura Ross

**Affiliations:** Institute of Evolutionary Biology, University of Edinburgh, Edinburgh, EH9 3JT, UK; Department of Genetics, Evolution and Environment, University College London, London, Gower Street, London, WC1E 6BT; Whitehead Institute for Biomedical Research and Department of Biology, Massachusetts Institute of Technology, Howard Hughes Medical Institute, Cambridge, MA, USA

**Keywords:** paternal genome elimination, mate preferences, sex ratio bias, sperm limitation, male fitness, sexual conflict, Diptera: Sciaridae, Sciara coprophila

## Abstract

Mate quality and the cost of mating affect the evolution of mating preferences and is one reason females often show stronger mate preferences than males. Fungus gnats in the family Sciaridae (Diptera) are a family in which we might expect to see the evolution of strong male mate preferences. Many Sciaridae species are monogenic, where females exclusively produce offspring of one sex. Sciaridae species also exhibit paternal genome elimination, a reproductive system where males only transmit maternally inherited chromosomes to offspring. Therefore, Sciaridae males would benefit from exhibiting mating preferences for females that produce female offspring, as a male’s genes are only transmitted to future generations through his daughters, not his sons. We explore male mate choice in the sciarid fly *Bradysia* (formerly *Sciara*) *coprophila*. We find that mating is costly, as males become sperm limited through multiple matings, and that males exhibit preferences for larger females, suggesting that males are selected to be choosy. However, we do not find male preferences for females that produce female offspring, instead we find that males prefer mating with females that produce male offspring. We speculate that this seemingly maladaptive behaviour may be due to female receptivity rather than male preference, or that males are unable to distinguish between females of different types, which is perhaps surprising since these females differ genetically by 1000s of genes (through a large paracentric inversion on the X chromosome). Together we show how the interplay between unusual genetics and sex determining systems may affect mating system evolution.

**Summary statement:** In the fungus gnat *Bradysia coprophila* females are genetically predetermined to produce broods of just one sex and males only transmit maternally inherited genes to offspring. These factors suggest males should have strong mating preferences for females that produce daughters, which we explore. We find that while males would benefit from being “choosy”, they appear unable to distinguish the two female types, possibly because females are selected to hide their sex determining phenotype.

## Introduction

Reproductive success of both males and females depends on the fecundity, fitness, and parental investment of their partners. Mate choice decisions can therefore have significant fitness implications. Mating preferences are predicted to evolve whenever benefits arising from variation in mate quality, or costs imposed through mating, favour discrimination between mates, and the level of choosiness displayed by each sex should reflect the contribution of that sex to the offspring (Trivers, 1972). In general, females are expected to exhibit stronger mating preferences than males because they are most often the sex for whom both parental investment and the costs of gamete production are greater (Servedio, 2007; Parker et al., 1972). As such, research on mating preferences often focuses on understanding female choice (Gavrilets et al., 2001; reviewed by Kelly, 2018). Males also experience costs associated with mating such as ejaculate production, sperm depletion, allocation of time spent looking for mates, reduced opportunities to mate with other partners, and production of nuptial gifts (reviewed in Scharf et al. 2013). Furthermore, males are predicted to display mating preferences, even if their contribution to parental investment is negligible, when there is sufficient variance in female quality (Ryan et al., 1996; Edward and Chapman, 2011). There is evidence of male mate choice across many insect clades (reviewed in Bonduriansky, 2001). Males may discriminate among females that differ in traits that serve as reliable predictors of fecundity, such as age, mating status), and body size (Friberg, 2006; Xu and Wang, 2009; Lüpold et al., 2011; Cotton et al. 2015). However, sexual conflict can also play into mating dynamics, as mating involves two individuals with different genotypes, and the outcome that would result in the highest fitness for one individual may not be the same as for the other, which can lead to the evolution of various methods to manipulate mating interactions (Parker, 2006).

Another relatively unexplored factor that can influence mate quality is the presence of selfish genetic elements such as selfish chromosomes or meiotic drivers. One way that meiotic drive alleles can bias their representation is by killing gametes and causing sex ratio distortion. In such cases, preferentially mating with partners without drivers will allow an individual to make a greater genetic contribution to future generations (reviewed in Lindholm et al. 2016). For example, in stalk-eyed flies, the presence of a meiotic driver that kills Y-bearing sperm has been known to result in female preference for males that lack driving alleles (Presgraves et al., 1997; Wilkinson et al. 1999). A particularly unusual form of meiotic drive occurs in the fly family Sciaridae (black-winged fungus gnats). Sciaridae species exhibit a non-Mendelian chromosome transmission system, found in several arthropod lineages, known as paternal genome elimination (PGE) (Metz, 1938; reviewed in Gerbi, 2022). Under PGE, males only transmit maternally inherited chromosomes to offspring. In contrast, females transmit chromosomes to offspring in a typical Mendelian manner (i.e. the chromosome inherited from either parent has an equal probability of being transmitted into eggs). Sciaridae species also have an XO sex chromosome system (i.e. females have two X chromosomes and males have one) which is regulated through chromosome elimination early in embryogenesis (Du Bois, 1933).

Interestingly, females of many sciarid species are also monogenic, meaning that females only produce offspring of one sex, and are either gynogenic (produce only female offspring) or androgenic (produce only male offspring) (Metz, 1931). In the sciarid *Bradysia coprophila,* whether females produce female vs. male offspring is genetically determined by a large X-linked inversion (Metz and Schmuck, 1929; Crouse et al., 1977) with androgenic females carrying two typical X chromosomes, and gynogenic females carrying one typical X chromosome and one X chromosome with a large, paracentric inversion (Metz, 1931, 1938; Crouse et al., 1977; Baird et al., 2023).

We would expect Sciaridae males to exhibit mating preferences for gynogenic females. Androgenic females only produce sons, which due to PGE only transmit their mothers’ chromosomes to their offspring. Therefore, males gain no fitness through mating with androgenic females (Figure 1). In contrast, gynogenic females produce female offspring which will transmit their father’s genes to future generations. However, male mating preference against androgenic females at a population level could have important consequences, as male preferences towards gynogenic females would skew the population sex ratio towards females.

**Figure 1.**
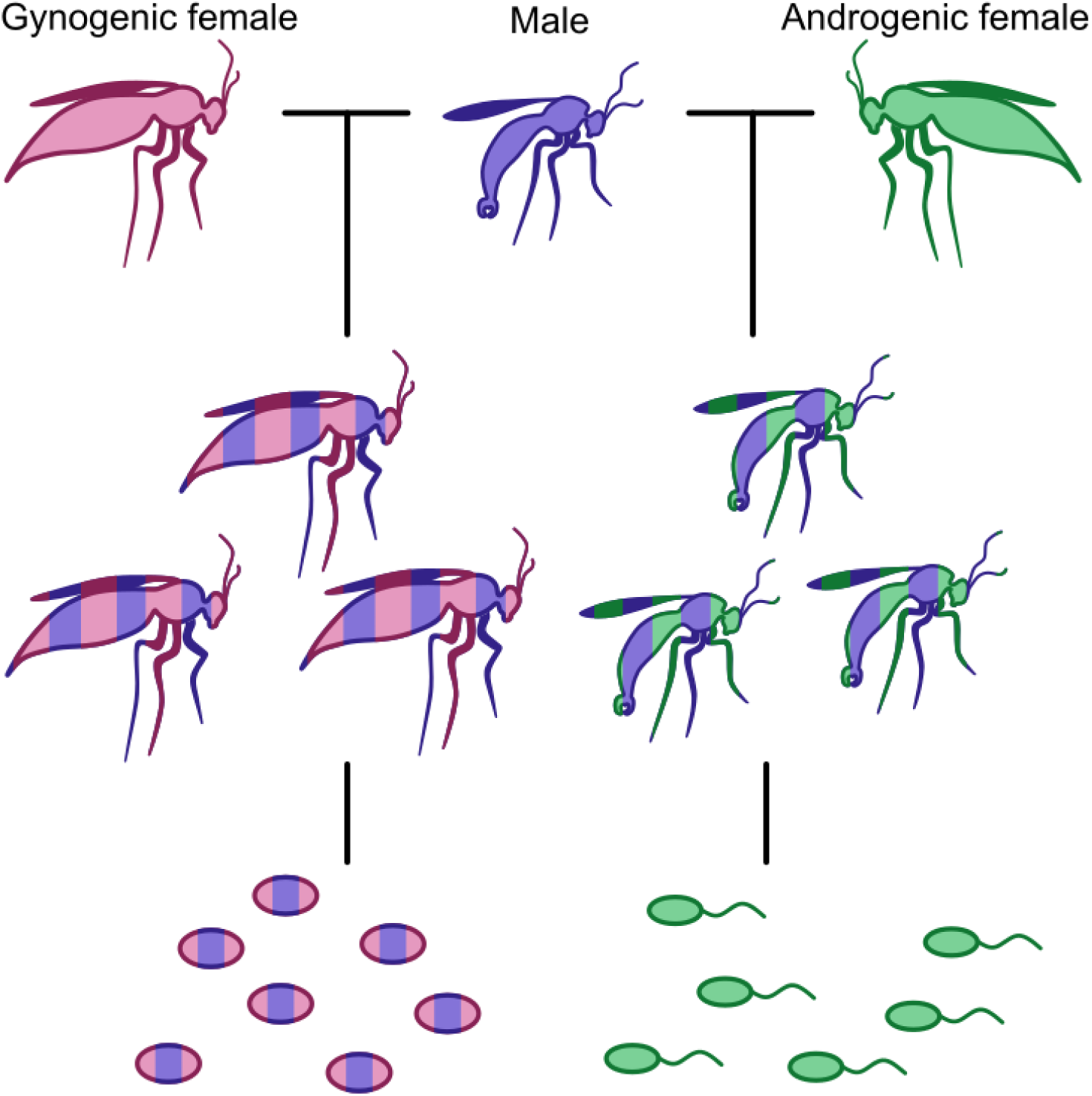
The reproductive system in the sciarid *Bradysia coprophila*. Females are monogenic and produce either female (pink female-gynogenic) or male (green female-androgenic) offspring only. Both male and female offspring carry genes transmitted from both their mother and father (yellow), but male offspring only transmit maternally inherited genes through sperm (i.e. sperm only contain genes from the androgenic female), while female offspring transmit genes in a Mendelian manner (i.e. eggs contain both male and gynogenic female genes). Therefore, the male will only gain fitness from producing female offspring.

Additionally, as there is never a benefit to males for mating with androgenic females, we might expect androgenic females to be under strong selection to not advertise their female type to males. Androgenic and gynogenic females differ by a large segregating X-linked inversion, with extensive divergence in gene expression between the two female morphs (Baird et al. 2023), which may present opportunities for females to diverge in their behavior or, conversely, for males to identify females based on divergent phenotypic differences.

Flies in the Sciaridae genus *Bradysia* have been extensively studied in a lab setting (Gerbi 2022), but little is known about their biology in natural populations. This is important as the development of mate preferences depends not only on genetic but on ecological and biological factors, such as whether mating is costly for males, and how many opportunities a male has to mate (Bonduriansky, 2001). *Bradysia* are known to exhibit elaborate courtship rituals where males display courtship behaviours such as wing flicking and abdominal thrusts towards females to initiate a mating attempt. Females can resist male mating attempts either through moving away from a courting male or kicking the male (Liu et al., 2002; Featherston et al., 2013). Elaborate or lengthy courtship behaviours generally occur in species in which male competition for females is strong but may impose costs on males through investment in time and energy, and reduction in future mating opportunities (Bonduriansky, 2001). Furthermore, *Bradysia* adults live only a few days as an adult, with females only mating once and producing a single clutch of eggs soon after mating (Moses & Metz 1928). Males also only undergo meiosis in early pupal development, and so have a finite amount of sperm as an adult (Mattingly & Dumont 1971). Together, these factors indicate that mating may be costly for male *B. coprophila* and selection may therefore favour the evolution of male mating preferences in this species.

However, previous work found that females exhibit mate preferences such that gynogenic females resist mating attempts more often than androgenic females (Featherston et al., 2013). This is in line with expectations: the offspring of androgenic females will not transmit their father’s genes to their progeny, so there is likely a lower selection pressure for choosiness in androgenic compared to gynogenic females. The expectations for mate preferences in Sciaridae therefore depend on the cost imposed on males when mating with androgenic females, the ability of males to recognise the two female morphs, the capacity for females to avoid recognition by males, and differences in female resistance behaviour.

In this study we conducted experiments on a laboratory line and a recently collected line of *B. coprophila* to understand (i) if males experience costs due to mating, and (ii) whether males exhibit mate preferences as a result of PGE and monogeny. First we examined whether males were limited by the number of females they could mate with and the extent of sperm depletion after mating. Secondly, we assessed the ability and willingness of males to choose females. We altered female quality (which we manipulate by raising females at different densities), to first assess if males had the capacity to choose based on a reliable predictor of fecundity (Bonduriansky, 2001; Lüpold et al., 2011). We then performed male mate choice assays for female type, to see if males could distinguish between androgenic and gynogenic females. We find that mating does appear to be costly for males as they are limited by both the number of females they can mate with and the amount of sperm that they have. Males also show mate preferences based on both female quality and female type. However, male mate preferences are weak and, surprisingly, we find that males prefer to mate with androgenic rather than gynogenic females. We hypothesize that this seemingly counterintuitive behaviour is a result of divergence in female receptivity rather than male mate choice, and that this could be due to stronger choosiness by gynogenic females (Featherston et al. 2013), or a strong evolutionary response by androgenic females to secure a mate. Alternatively, the biology of this species may mean that the benefit to evolving mate preferences for males is slight compared to the cost of missing a mating opportunity. Therefore, we suggest that sexual antagonism, perhaps combined with weak selection on male mating behaviour, may be shaping mating dynamics in *B. coprophila*.

## Methods

### Culture information

We conducted mating experiments on two *Bradysia coprophila* lines. The lab line has been kept in laboratory conditions since the early 20th century (Metz & Smith 1931), while the other line was collected from house plants in Edinburgh in 2018 by C. Hodson and maintained in the lab since then, therefore, we refer to it as the CH line. We maintain both fly lines by mating individual females with one or two males in a glass vial (25mm diameter x 95mm) with bacteriological agar, leave the female to lay eggs and raise the larvae by sprinkling a powdered food mixture (of organic wheat straw, mushroom powder, spinach powder and brewers yeast) into the vial every 2-3 days until the larvae pupate. Female *B. coprophila* from the lab line have a wing polymorphism on the X chromosome which causes females that carry it to have wavy wings (Metz & Smith, 1931). As this wing polymorphism is a dominant marker only present in females that produce female offspring, it can be used to identify gynogenic from androgenic females. This polymorphism is not present in the CH line. However, for both the lab line and the CH line, it is important to note that physical differences between gynogenic and androgenic females have not been examined, and there may be traits important for mating success that differ between the female types.

### Male mating capacity experiment

In order to gather a more complete understanding of male mating behaviour, we conducted a mate choice experiment to determine whether males are limited in their mating capacity, whether they show preferences towards females of a certain type (i.e. androgenic or gynogenic) when presented with both types of females at the same time, and how offspring production differs by female type. We conducted this experiment with both the lab fly line and the CH fly line. We predominantly used the lab line, as we are easily able to distinguish androgenic from gynogenic females in this line. However, we also wanted to compare the lab line to a recently collected CH fly line, to determine whether lengthy lab history of this line affected their mating behaviour. For the trials with the lab line, we raised males at different densities (either ∼10 or ∼50 larvae per vial) to produce males of different sizes (Pitnick, 1991; Byrne and Rice, 2006). Males raised at different densities are expected to be different sizes as adults due to different levels of crowding, which we expected to result in males having different amounts of energy reserves. For the trials with the CH flies, we conducted experiments with male flies raised only at high densities. All females used in this experiment were raised in vials of approximately 50 larvae. For each experimental condition (high density lab males, low density lab males, high density CH males) we conducted between 22-24 replicates.

To prepare the mating vial, we added 10 females into the vial from the same fly line as the male (Figure 2). For trials from the lab line, the mating vial included five androgenic and five gynogenic females (0-4 days old, measured as time since eclosion) whereas the CH fly line contained 10 random 0-4 day old females. We determined female type for flies from the CH line once they produced offspring (i.e. we could only determine the female type of mated females). We added a 0-4 day old male to the mating vial with the 10 females. We left the mating vial for 24 hours, then removed and froze the male, and moved each female into a separate vial (9cm x 2.5 diameter) filled with approximately 2 cm of agar. We took note of any females that died during the trial and left the remaining females to oviposit. We removed and froze the female once she died and noted whether she produced eggs. We left the vials for 14 days from the day the mating trial was started (by which time we expect all viable eggs to have hatched), then counted the larvae, taking the average of two counts from each vial. We recorded the number of females in each mating trial that produced larvae and the number of larvae that each female produced. For flies from the lab line we also noted which type of female produced offspring while for the CH fly line we determined the female type by raising the larvae until adults and determining the sex of the first ten offspring. We measured the size of each male after the experiment by photographing the male with a calibration slide (with a 1 cm ruler divided into 0.1 mm increments) under a dissecting microscope fitted with a camera, then measuring the thorax width, as a proxy for body size, using FIJI (Schindelin et al., 2012) (Supplementary Figure 1).

**Figure 2.**
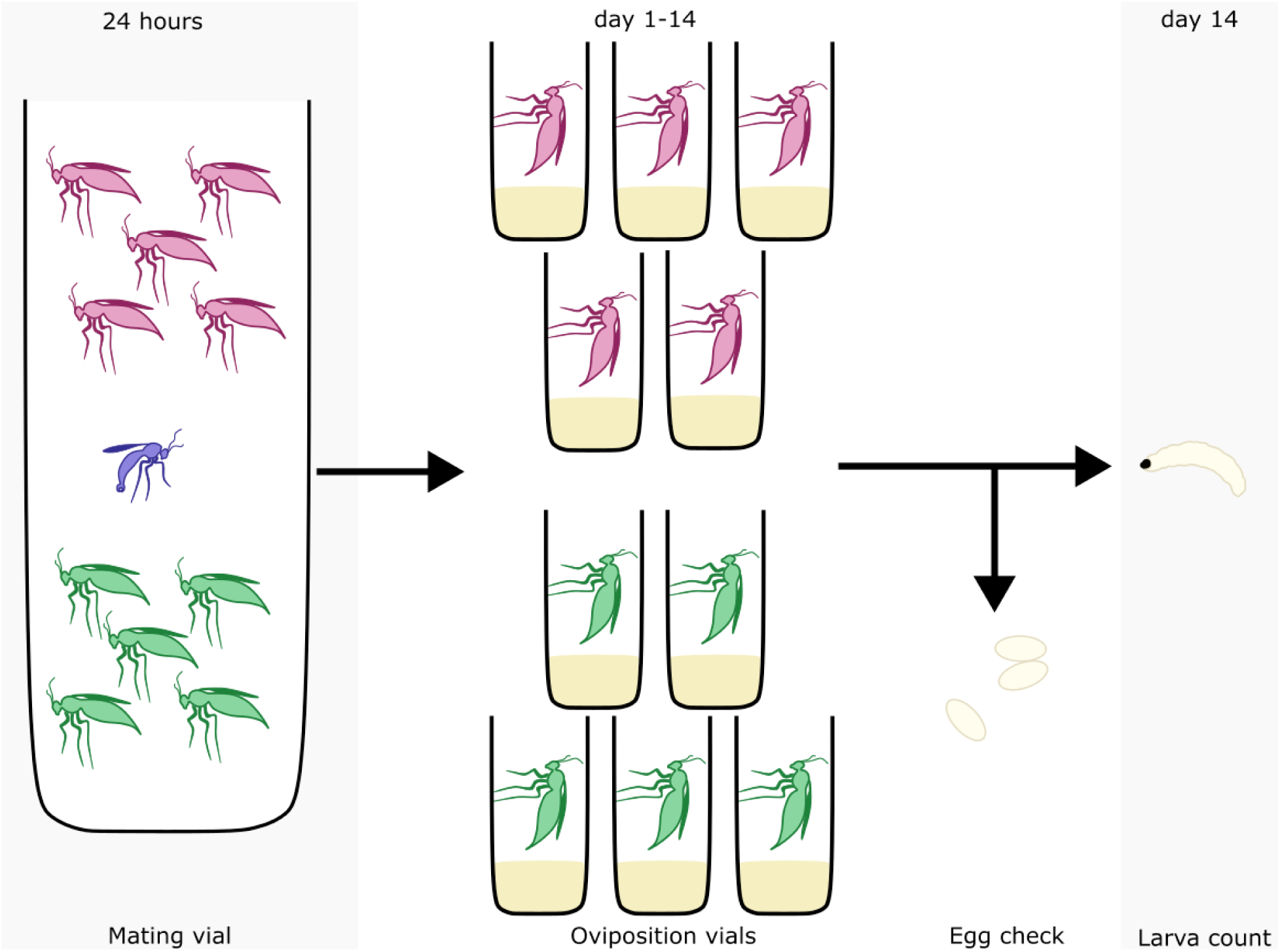
Schematic of mate capacity experiment. One male was housed with 10 females for 24 hours (5 gynogenic (pink-top) and 5 androgenic (green-bottom) for the lab line and 10 females of unknown type for the CH line). Afterwards, females were kept in separate vials for 14 days and we recorded whether they produced eggs. After 14 days, the larvae each female produced were counted.

We also measured the thorax size of a subset of the females in the experiment (all 10 females from 16 trials with the lab line and 15 trials with the CH line) to determine whether female size correlates with the number of offspring she produces.

For the mating trials with the lab fly line, we also measured the number of sperm in the testes of 11 males from each condition after the mating trial. We dissected the testes out of these males in a drop of 1X PBS by gently pulling on the males’ genital claspers with tweezers while holding the abdomen until the testes came apart from the body. We separated the testes with insect pins, then spread the sperm around the slide. We stained the sperm by fixing the tissue in 5µl of 45% acetic acid. We allowed the slide to almost dry, then added a drop of DAPI with Vectashield (Vector laboratories), and placed a coverslip on the slide, sealing the edges with nail polish. We viewed the slides under a fluorescent microscope with a DAPI filter (Leica DM 2000 LED microscope) and counted the number of sperm present on each slide. In a subset of counts (n=16), we found that males had either very few sperm after the mating trials (<150) or many sperm, and that males that had many sperm often had more than 1000 (n=6) (Supplementary Table 1). We chose to count to a maximum of 200 sperm for analyses as counting to 1000 sperm was time prohibitive.

### Male mating preference experiment

We conducted an experiment to examine male mating behaviour towards females of different types and sizes. For this experiment, we used flies from the lab line and made vials with female larvae raised at high densities (∼50) or low densities (∼10), to produce smaller and larger females respectively. We conducted consecutive mating trials in which one 0-4 day old male was allowed to mate with 0-4 day old virgin females of a different condition (low/high density upbringing) and type (gynogenic/androgenic), with the four mating trials for each female type x size randomized and occurring over one day. We conducted mating trials with 22 males in total (i.e. 88 behavioural observations).

On the day of the behavioural trial, we transferred the male into a plastic vial ¾ full of agar (to limit the amount of space in the vial and more easily view the trial under the microscope). We let the male acclimatize for 5 minutes and placed a female from one of the four conditions in a randomly generated order into the mating arena with the male. We then began recording the behaviour of the male and female under a dissecting microscope (Leica dissecting microscope with an attached MC170 HD lens). We recorded until the pair finished mating or for 20 minutes if mating did not occur. We then removed the female from the mating arena and froze her at-20°C. We left the male for 45 minutes, then conducted another mating trial with a female from one of the other conditions. The male was given the opportunity to mate with a female from all four conditions over the experiment day, then was frozen afterwards. We later viewed the recordings and noted whether mating occurred, when mating started and finished, and characteristic mating behaviours displayed by both the male and female (Liu et al., 2002; Featherston et al., 2013). To initiate mating, *Bradysia* males will first display a wing flicking behaviour towards the female, then will curl his abdomen and thrust towards the female.

Following this, the male will attach himself to the female’s abdomen with his claspers and adjust himself so he is positioned correctly to inseminate the female, after which copulation will occur (Supplementary video 1). At any time during this sequence, the female may display resistance behaviour in the form of either walking away from the male or kicking the male away. When scoring the mating trials, we recorded whether the male exhibited the wing flick behaviour, the number of abdomen thrusts the male made towards the female, the number of times the male attached to the female, whether the female exhibited resistance behaviour in response to mating attempts made by the male, and whether the male or female terminated the mating act.

After each mating trial, we measured the size of all females in the same way as for the mating capacity experiment. We also estimated the number of sperm transferred to the spermatheca of females that had successfully mated to assess whether males exhibit cryptic mate choice through reducing sperm transferred to less desirable females (Wedell et al. 2002). To do this, we dissected the spermatheca out of each female using insect pins in a small drop of 1X PBS on a microscope slide, then stained the sperm in the spermatheca using the same method outlined above for the male reproductive tract.

### Data analysis

We carried out all statistical analyses using RStudio v.3.6.3 (R Core Team, 2020) using ggplot2 to produce figures (Wickham, 2016). We used package lme4 for generalised linear mixed models, and package nlme for linear mixed effects models (Bates et al., 2015; Pinheiro et al., 2020). We inferred p-values for fixed effects using the Satterthwaite approximation to calculate the degrees of freedom with the package lmerTest (Kuznetsova et al., 2017).

#### Male mating capacity experiment

For the trials with the lab line in which we manipulated male density, we determined whether male size differed between density treatments with a Welsh’s two-sided t-test with male density as the fixed effect and male thorax size as the response variable. We compared whether males mated with less than the available number of females for both the lab line and the CH fly line with a Welsh’s one-sided t-test. We censored any females from the analysis which we could not say whether they had mated either because they died or escaped during the mating trial. For all the mating capacity analyses, we scored females as having mated if they produced eggs which hatched into larvae. This is because females can produce unfertilized eggs which do not hatch if they have not mated. As we could not tell whether the females from the CH line were androgenic or gynogenic unless they produced offspring, we analysed whether males displayed mating preferences in a different way than for the lab line. For the flies from the lab line, we analysed whether males of different conditions mated with different numbers and types of females with a generalised linear mixed effect model with a binomial distribution. We used whether the female mated as the response variable (i.e. 1=mated, 0=unmated), and male condition treatment (either high or low density) and female type (gynogenic or androgenic), and an interaction between these two factors as fixed effects. We also included male ID as a random effect. For the trials with the CH fly line, we used a generalised linear model with a poisson distribution. For this model, we included the number of females in each trial of each type that mated as the dependent variable, and female type as the fixed effect.

To get an estimate of the total number of offspring (i.e. larvae) a male produces, and to explore whether the number of offspring, and the type of females a male produces offspring with, is dependent on his condition, we used a generalised linear mixed effect model with a poisson distribution. We rounded the count of the number of larvae each male produced of each sex to the nearest integer and used this value as the response variable, and included female type, male density, and an interaction between these factors as fixed effects, and male ID and observation as random effects. We used a similar model for the CH line but did not include male rearing density or the interaction term as fixed effects. We assessed whether female size influenced the number of offspring produced by determining for a subset of the trials in which we measured the female size as well as male size, whether female size influenced the number of offspring she produced. For the lab fly line and the CH fly line, we used a generalised linear mixed effect model with a poisson distribution, and the number of larvae produced by each female as the response variable. We included female thorax length and female type as fixed effects and male ID and observation as random effects. For the flies in the lab line, we also measured the number of sperm remaining in the testes for 22 males after the mating trial. We analysed whether sperm number was affected by the number of females the male had mated with a generalised linear model with a poisson distribution, with sperm number (to a maximum of 200 sperm) as the response variable, the number of mates as the fixed effect and observation as a random effect.

#### Male mating preference experiment

For the mating preference experiment, we determined whether female size differed between density treatments using a Welsh’s two-sided t-test with the female density as a fixed effect and female thorax size as the dependent variable. To determine whether males showed mating preferences we used a generalised linear mixed effect model with a binary distribution. We included whether mating occurred as the response variable (1=mated, 0=not mated), the mating order, density treatment (either high or low density), female type (gynogenic or androgenic), and an interaction between these two factors as fixed effects, and male ID as a random effect. We also analysed the amount of effort males will place into mating with different types of females, by conducting a generalised linear mixed effect model with a poisson distribution, with the number of times males thrust towards a female as the response variable and female type, density females were raised at, an interaction between these two factors, and mating order as fixed effects. We included male ID as a random effect and an observation level random effect to account for overdispersion (according to Harrison, 2014). We analysed whether copulation duration is affected by female type or density with a linear mixed effects model. We log transformed mating duration and used this value as the response variable and included the same fixed and random effects as the model above. To assess whether males transferred more sperm to certain types of females, we analysed sperm counts for mated females using a generalized linear mixed effects model with a poisson distribution, with sperm number as the response variable. We included density treatment, female type, an interaction between these two factors and mating order as fixed effects, and male as a random effect as well as an observation level random effect.

## Results

### Male mating capacity experiment

We raised male flies at low and high densities as a method to generate differences in the condition of males. Although we found that males raised at a low density were on average larger than males raised at a high density (average thorax size of 0.67 and 0.63 respectively) this difference was non-significant (Supplementary Figure 2) (t.test, df=19.06, p=0.07). For the mating capacity experiments with both the lab line and the CH line, all males and 98.14% of females survived for the 24 hours in the mating vial. The females that did not survive were excluded from analyses. In the 24 hours that males were left to mate, males in both conditions (from the lab line) and the CH line mated with fewer than the total number of available females (lab line-t.test, df= 42.35, p-value < 0.0001, CH line-t.test, df= 20.56, t=-11.65, p<0.0001). On average, males in the lab line mated with 4.85 of the available 9.93 females (note that the available number of females was slightly less than 10 as we censored females that died or escaped from the mating vial) while males from the CH line mated with on average 5.24 of the available 9.95 females.

We examined the probability that a female had mated based on whether she is gynogenic or androgenic and the condition of the male she had access to mate with. We found that male condition did not affect the likelihood that females mated (Supplementary Table 2; binary glmer: est:-0.12, s.e.= 32 z=-0.39, p=0.70), but that androgenic females were more likely to mate than gynogenic females (Supplementary Table 2; binary glmer: est: 0.56, s.e.=0.28, z=1.96, p=0.050) and there was no significant interaction between these two factors (Figure 3A). Therefore, males mated more often with androgenic rather than gynogenic females. However, unlike the lab line, males from the CH fly line did not show a preference for either gynogenic or androgenic females (Figure 3B) (Supplementary Table 3; glm: est=-0.07, s.e.= 0.19, z=-0.39. p=0.70)

**Figure 3.**
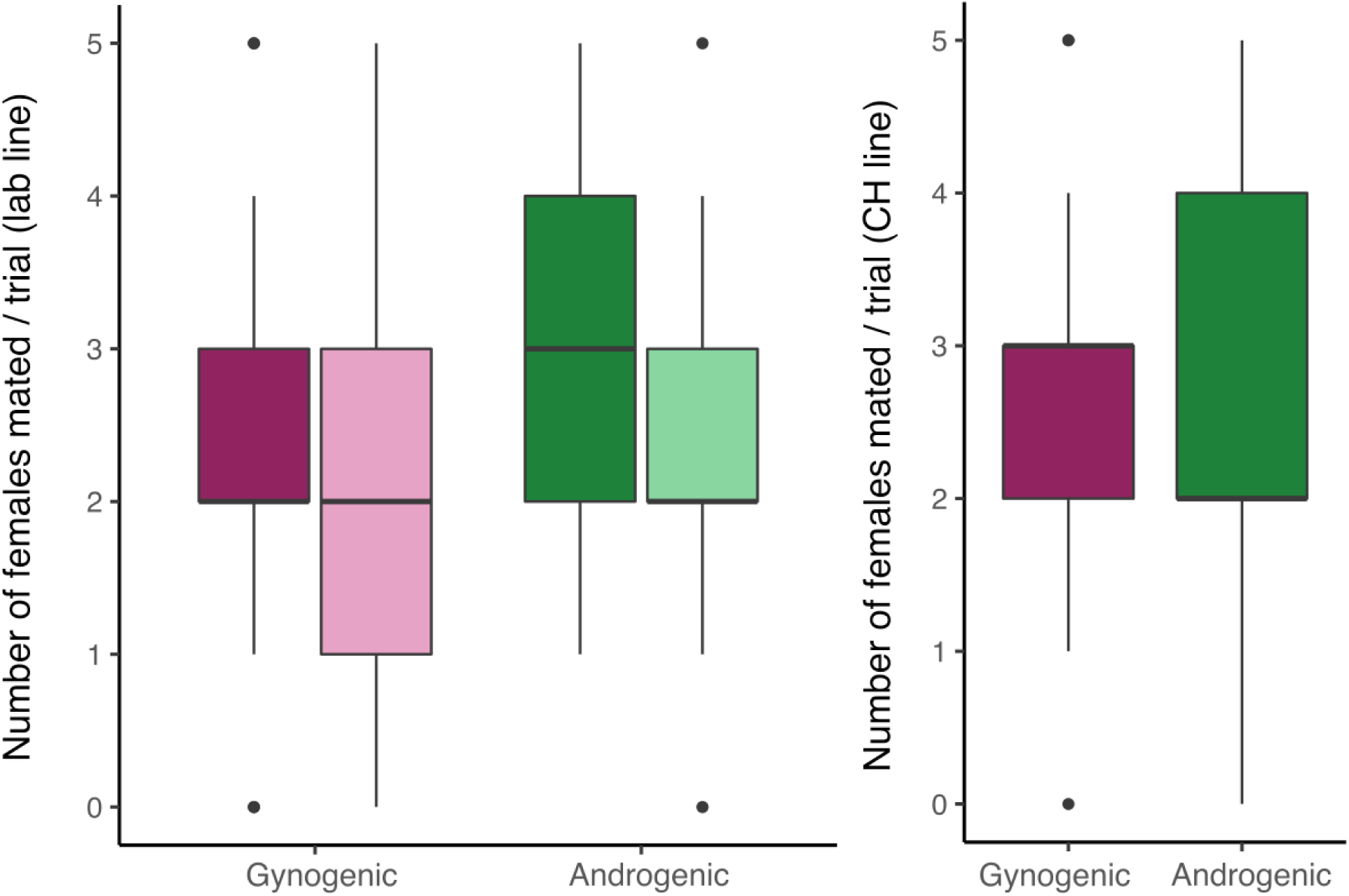
(A) Number of gynogenic (pink) and androgenic (green) females mated per trial. Androgenic females mate more often than gynogenic females. Trials with a low density male (N=20) are shown in a light colour (right side), and those with a high density male (N=22) are shown in a dark colour (left side). (B) Number of gynogenic (pink) and androgenic (green) females that males in the CH fly line mated with per trial (N=21). Males displayed no mating preferences towards either female type.

We examined whether the number of offspring produced by each male depended on female type or male condition. For the lab fly line, we found that males produced approximately the same number of male and female offspring (Supplementary Table 4, glmer: est=-0.34, s.e.= 0.27, z=-1.24, p=0.22), with males producing a mean of 123.3 male and 122.2 female offspring. Offspring number did not depend on the density a male was raised at (Supplementary Table 4, glmer: est=-0.63, s.e.= 0.40, z=-1.60, p=0.11). Additionally, there was no interaction between these two factors (Figure 4A). For the CH fly line, males also produced approximately the same number of male and female offspring (Figure 4B; Supplementary Table 5; glmer: estimate: - 0.26, s.e.= 0.40, z=-0.64, p=0.52), with males producing a mean of 155.8 female larvae and 145.4 male larvae. We also examined for a subset of trials whether female size influences the number of offspring she produced. We found that larger females produced more offspring than smaller females in both the lab fly line (Figure 4C) (Supplementary Table 6; glmer: est= 22.17, s.e.= 7.13, z=3.11, p<0.01) and CH fly line (Figure 4D) (Supplementary Table 7; glmer: est= 2.43, sterr= 1.19, z=2.03, p=0.04) showing that female size is indeed an indicator of fecundity. However, female type was not a predictor of offspring number for either fly line (Supplementary Table 6, 7).

**Figure 4.**
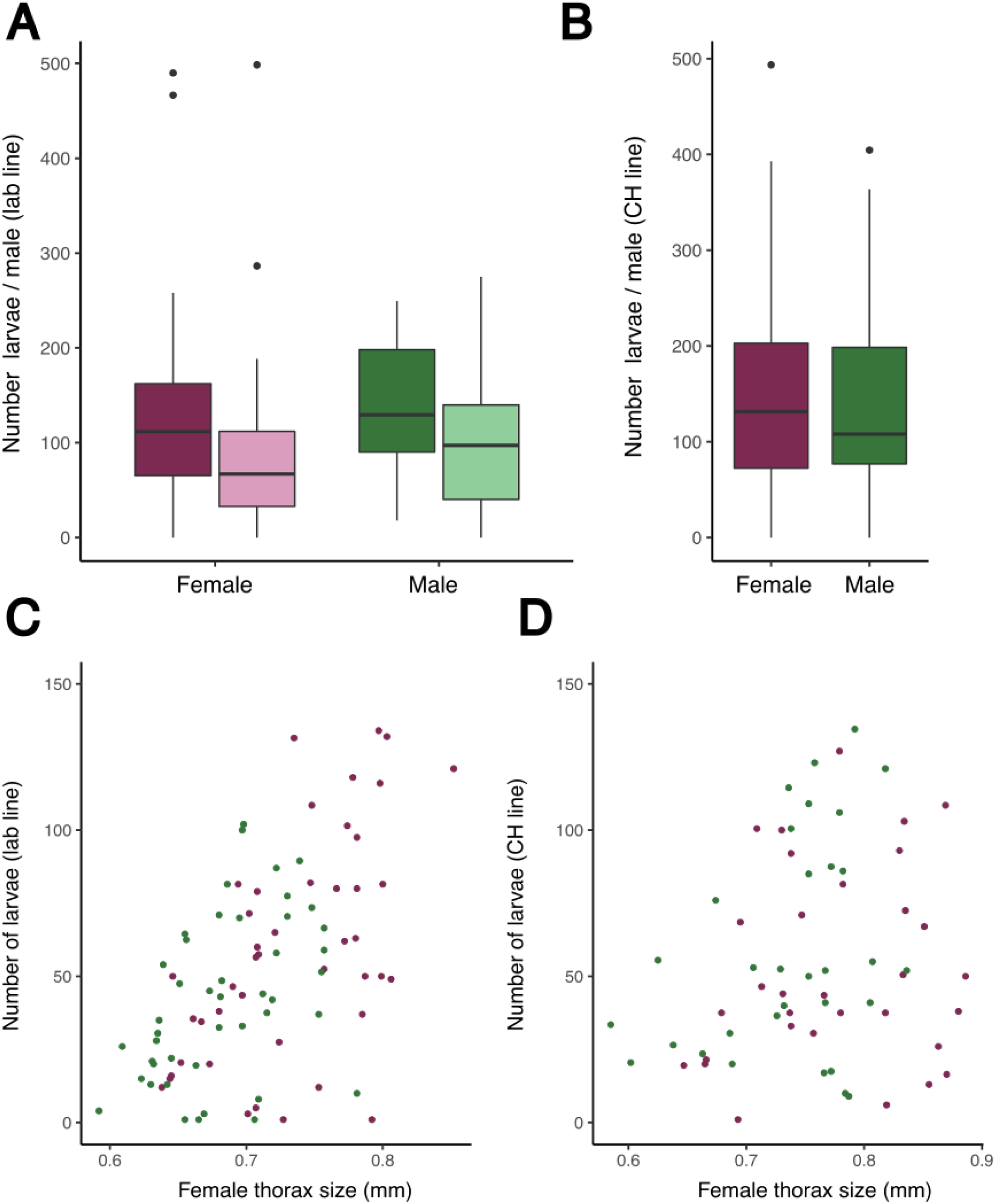
(A) Number of female and male larvae produced by males in the high density (dark colour-left side) and low density (light colour-right side) treatments from the lab line. (B) Number of female (pink) and male (green) larvae produced by males from the CH line. Number of larvae produced by females from the lab fly line (C) (n=84), and the CH fly line (D) (N=64) based on female thorax size. For females from both the lab and CH *B. coprophila* lines, larger females produce more larvae than smaller females. gynogenic females=pink, androgenic= green.

We measured the amount of sperm in the testes of a subset of males after the mating experiment. We found that males that mated with more females had less sperm remaining in their testes compared to males that mated with fewer females, suggesting males can become sperm limited (Figure 5) (Supplementary table 8; glmer: est=-0.39, s.e.=0.19, z=-2.05, p=0.04). In fact, 10 males had very low levels of sperm after mating (50 or less).

**Figure 5.**
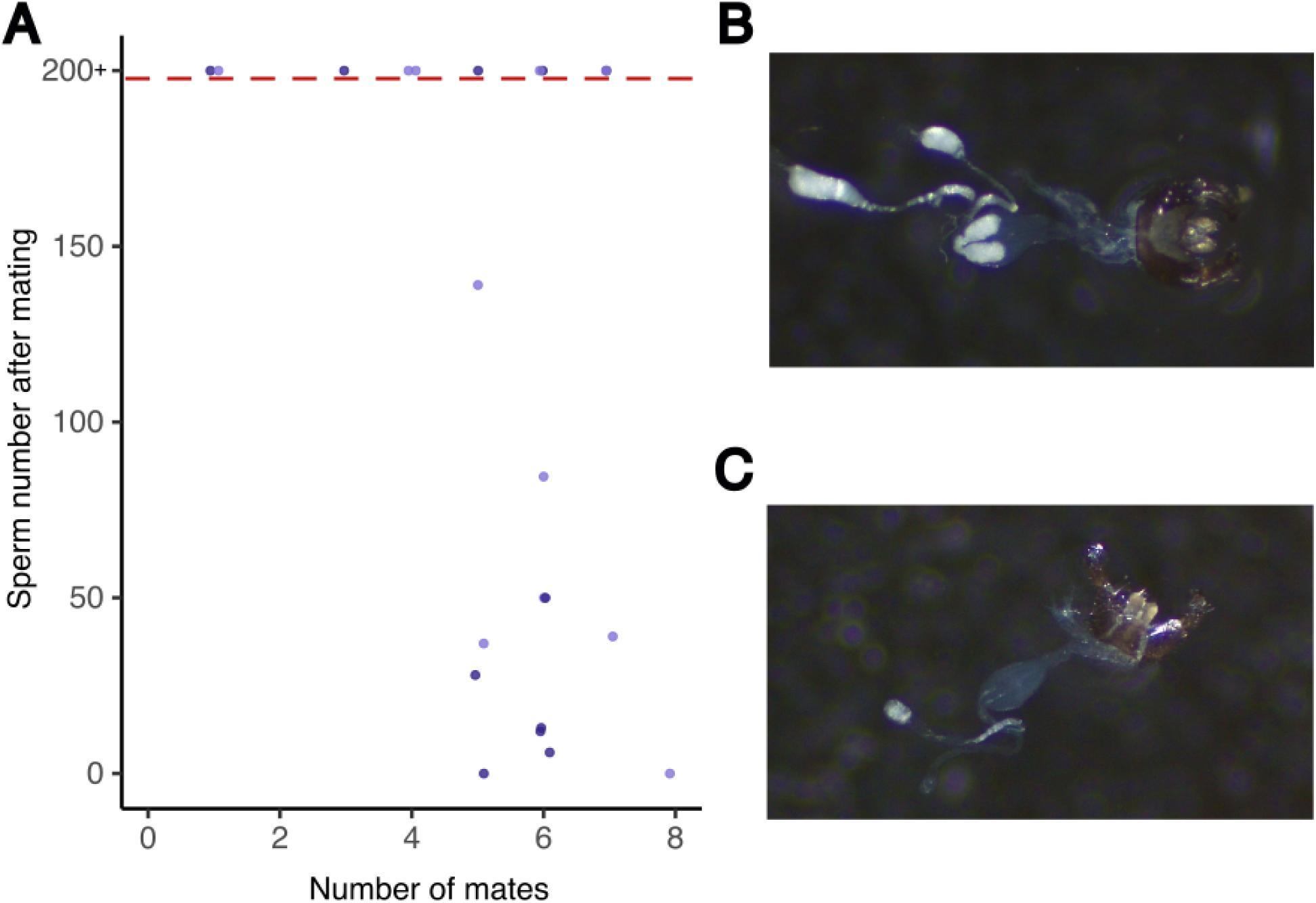
(A) Number of sperm in the testes of males (lab line n=22) after the male mate capacity experiment compared to the number of mates for each male. Males from the high density treatment are dark coloured and males from the low density treatment are light coloured. Data points are jittered (by 10% within each integer bin on the x-axis). The number of sperm was counted to a maximum of 200, the data points above the dotted line may have more than 200 sperm. (B-C) Representative images of male testes (attached to the claspers of the male) after the male mate capacity experiment. The male in (B) had more than 200 sperm in his testes after the mating trial (white mass in testes) but the male in (C) had only 13 sperm after the mating trial.

### Male mate preference experiment

To produce females of different body sizes we raised females at low and high densities. On emergence, females from high density treatment were smaller than females from low density treatments (t.test, df= 81.38, p<0.0001) (Supplementary Figure 3). Males in the mate preference experiment were more likely to mate with androgenic females than gynogenic females (Figure 6A; Supplementary Table 9: glmer: est=1.920, sterr=0.83, z=2.32, p=0.02) and females from the low density treatment (Supplementary Table 9: glmer: est=1.72, sterr=0.82, z=2.11, p=0.04) but there was no interaction between these two factors. Male mating order did not affect the probability of mating (Supplementary Table 9).

**Figure 6.**
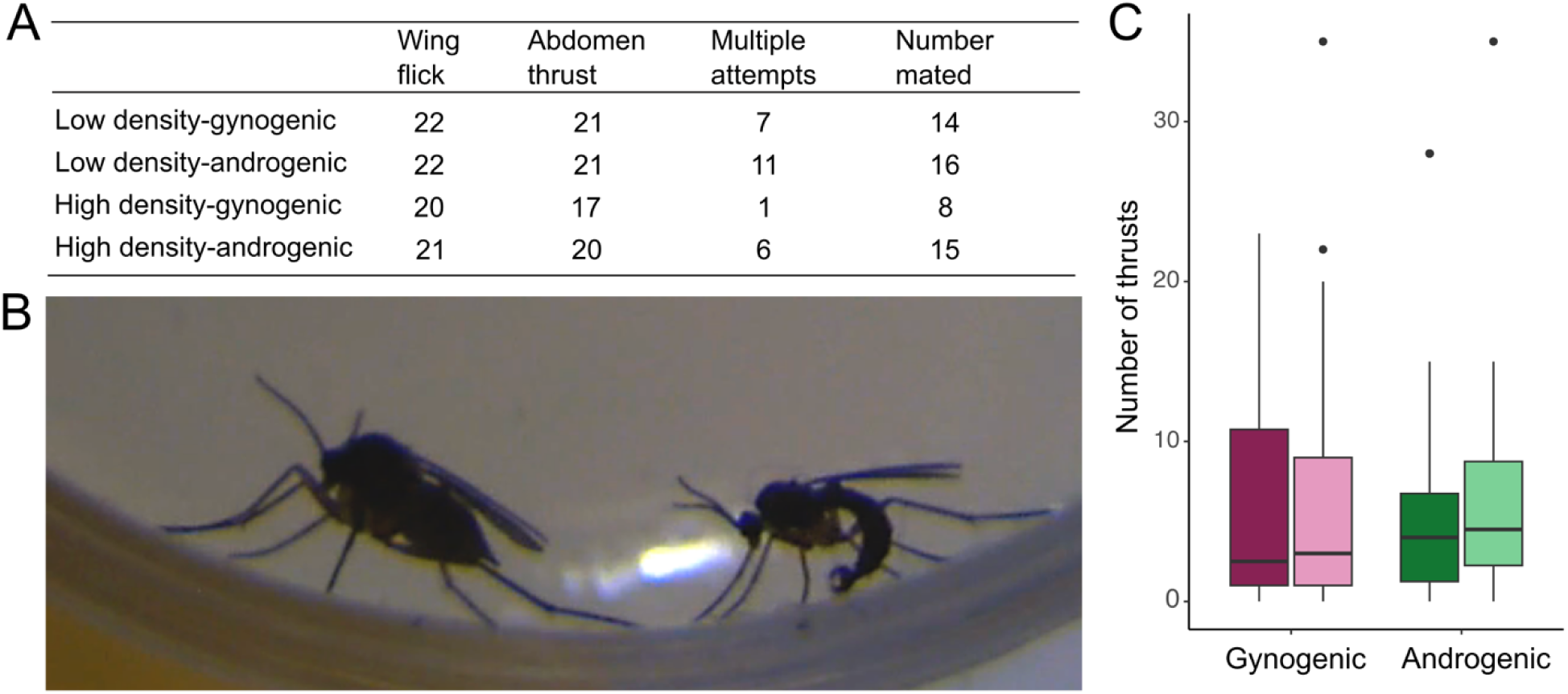
Mating behaviour of *B. coprophila* males. (A) Number of males that engaged in wing flick behaviour, abdominal thrust, and tried to mate with a female more than once for each type of female. The final column shows the number of females who mated in each category. Each male (n=22) was given an opportunity to mate with a female from each type. (B) An image of abdomen thrust behaviour where a male (right) tries to grab the female abdomen (left) in a mating attempt. (C) The number of abdominal thrusts (i.e. mating attempt) males directed towards each type of female.

To look at how individual behaviour affects whether a male ultimately mates with a female, we measured male and female behaviour during trails. In the vast majority of trials, males exhibited mating behavior in the form of wing flicks (96% of trials) and abdomen thrusts towards the female (90% of trials) (Figure 6A). In some instances, male mating attempts failed because of female resistance behaviour (29 trials), whereas in others males failed without female resistance, for example by falling off after clasping the female or abdomen curling when a female was too far away (13 trials). We found that the probability of a female rejecting a male’s mating attempt at least once was not dependent on female type or density (Supplementary Table 10). Additionally, the number of thrusts males exhibited towards each female did not depend on whether the male’s mating partner was gynogenic or androgenic or female size (Figure 6C; Supplementary Table 11).

Finally, we examined cryptic mate choice through mating duration and sperm transfer to females. Mating order was the most important predictor of mating duration, with males mating longer with females when they had been presented with more mating partners (Figure 7A; Supplementary Table 12; lmer: est=0.07, sterr=0.01, df=26.53, t=6.02, p<0.0001). There was also an interaction between female type and female density treatment (Figure 7B) (Supplementary Table 12; lmer: est=-0.11, sterr=0.06, df=27.05, t=-2.13, p=0.04). The amount of sperm females received through mating did not depend on either female type or female density treatment (Figure 7C), mating duration or mating order (Supplementary Table 13).

**Figure 7.**
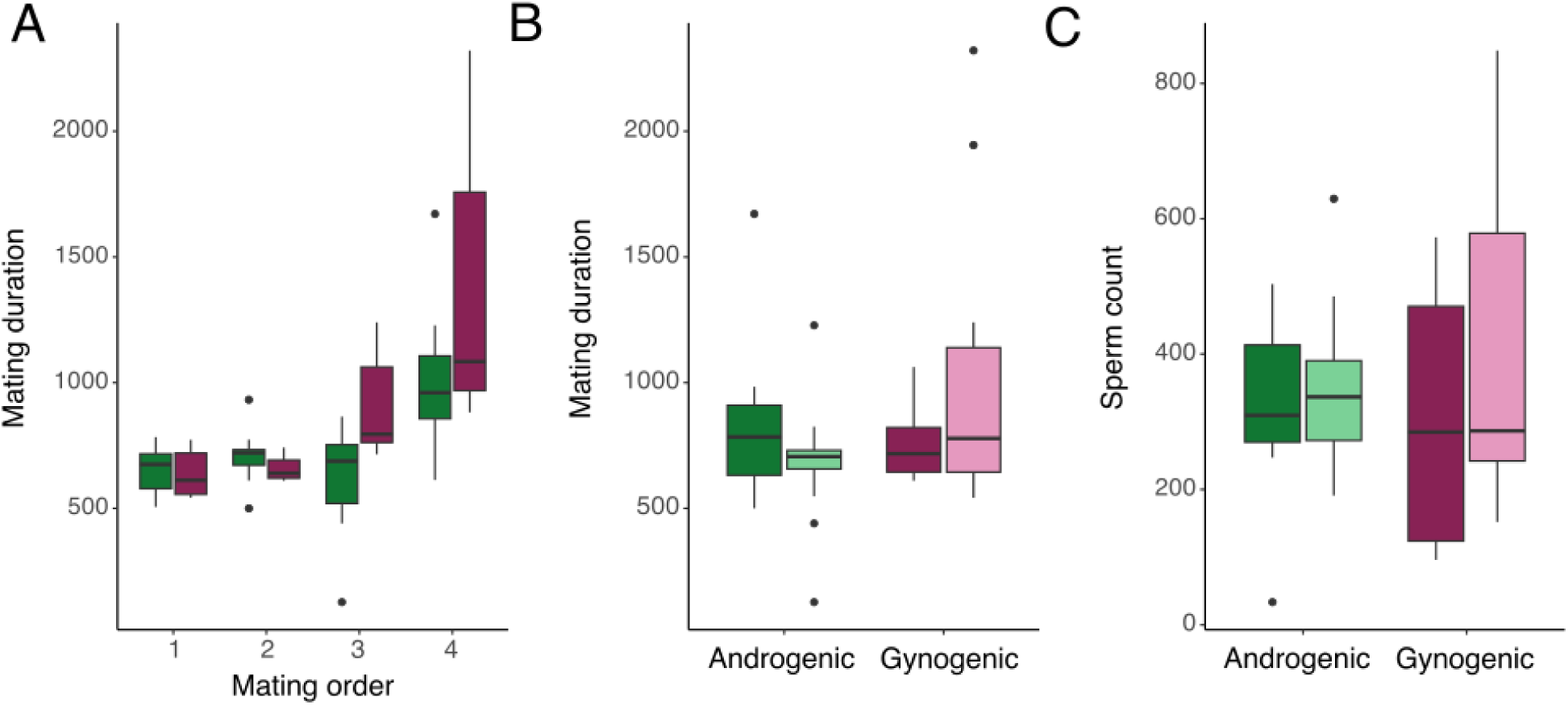
(A) Mating duration depending on the order the males were presented with females. Males took longer to mate with females over successive matings (gynogenic females-dark pink, androgenic females-dark green). (B) Mating duration (seconds) for copulations between males and androgenic (green) and gynogenic (pink) females raised at high (dark) or low (light) density (n=53). Mating duration did not depend on female type, or the density treatment (C) Number of sperm delivered by males to androgenic and gynogenic females of different density treatments (n=27). The amount of sperm males delivered to females did not depend on female type or density treatment.

## Discussion

The unusual inheritance system in Sciaridae leads to the prediction that males should be under strong selection to evolve mating preferences based on female genotype. Specifically, males should prefer to mate with gynogenic (female-producing) females, as these are the only females that will transmit their genes to future generations (see Figure 1). We conducted several experiments on *B. coprophila* to test this prediction, as well as to gather a better understanding of mating dynamics in this species, including how costly mating is for males and how female and male size affect mating dynamics.

We found that males have the capacity to exhibit mating preferences. Consistent with previous studies on male mate choice in other insects (Bonduriansky 2001), we find that males prefer to mate with larger females; a factor that we show is correlated with female reproductive output. Contrary to our expectations however, we found in both our mating behaviour experiment and our male mating capacity experiment, that males of our lab line of *B. coprophila* prefer mating with androgenic (male-producing) rather than gynogenic females. We found that males mated more frequently with these females in behavioural experiments, and when left with both androgenic and gynogenic females over a 24hr period, more androgenic females were mated at the end of this period. It is possible that for the lab line, the long history of laboratory rearing influenced this seemingly maladaptive behaviour, as male flies from the lab line are generally given access to only one female to mate with, obliviating selection for male choice.

We do not see this behaviour in the CH fly line, in which males did not display a preference towards mating with gynogenic or androgenic females.

The main question arising from our study is why do males in this system not show strong mating preferences, as predicted? We believe there may be several important factors informing this observation. One goal of this study was to understand more about the mating behaviour of *B. coprophila*, as we did not know how costly mating is for males, how likely males are to encounter multiple females in the wild, and many other factors that might affect actual mating decisions in wild populations. We found that mating for males does seem to be costly, as in a 24-hour period, males given access to 10 females were unable to mate with all of them and often had very low sperm reserves after the mating trials (approximately 50% of the males we measured had fewer than 100 sperm after the mating trial). As males do not continually produce sperm throughout their adult life (they produce it once as larvae), that effectively means that males have a finite number of females they can mate with. However, we do not know how many females a male is likely to encounter in more natural settings, and males were able to mate with up to 8 (and on average ∼5) females. It is possible that sperm limitation is not an important factor for mating decisions in natural populations, since males may rarely encounter more than 5 females in their adult life (although note that males did show a preference for large females in our experiment). In contrast, we would expect that mating in this species is quite costly for females, as they generally only mate once, and produce all their eggs once soon after mating (Moses and Metz, 1928), meaning that they have one opportunity to choose a mate. Given this, it makes sense that females, and specifically gynogenic females, are choosy (Featherston et al., 2013).

Our behavioural experiment also illuminated several important facets of mating behaviour in *B. coprophila*. We found that upon encountering a female, nearly all males would begin courting the female, exhibiting wing flick behaviour followed by abdominal thrusting towards the female. Additionally, males transferred a similar amount of sperm to females regardless of their type or condition. These observations add to the suggestion that males exhibit weak/no mate preferences and will attempt mating with almost any female they encounter. However, only about 60% of trials ended with a successful mating. Unlike the results from Featherston et al. (2013), we were unable to find evidence for differences in female resistance behaviour towards males between different female types, although our experiments were conducted in a slightly different setting and likely using a different species of *Bradysia*.

However, females did exhibit resistance behaviour in a sizable proportion of trials, and in 22 of the trials where mating did not occur, females exhibited resistance behaviour. This is suggestive that female behaviour might be driving some of the mating dynamics we witnessed. We focused primarily on male mating behaviour in this study, but we believe also taking female mating behaviour into account would help illuminate why males do not exhibit strong mate preferences. For instance, the surprising finding that males preferentially mate with androgenic females might be a consequence of sexually antagonistic interactions, as androgenic females need to secure a mating partner and so may be under strong selection to hide their type from males. Thus, differences in female receptivity between androgenic females, who need to secure a mating partner, and gynogenic females, who need to secure a *good* mating partner, may influence our results.

We also assessed whether males exhibited cryptic mate choice by measuring how much sperm males transferred to females in the mating behaviour experiment and the number of female and male offspring males produced in the male mating capacity experiment. In the behaviour experiment, males were the most likely partner to end copulation (in 77% of trials) and the number of previous mating partners for the male influenced mating duration, suggesting that males have some control over mating duration. One possibility for why males do not exhibit cryptic mate choice is that the dynamics in this system makes cryptic mate choice unnecessary. Since females generally mate only once, sperm competition is non-existent. As such, males may be selected to deliver as little sperm to their partner as will fertilize all her eggs (Parker & Pizzari, 2010). Therefore, in *B. coprophila*, selection for limiting sperm transfer to females, rather than mate preference, may be the factor driving male sperm allocation decisions. We are still untangling the factors that are important in *B. coprophila* mating dynamics, but our experiments, combined with previous work, suggest that sexually antagonistic interactions, along with the need to secure a mating partner (or partners for males), might be key in this system, and should be explored more thoroughly in future studies.

In other systems with meiotic drive, strong mate preferences have evolved against individuals that have drive alleles, both through avoiding mating with individuals carrying drivers and through cryptic mate choice (Angelard et al. 2008; Cotton et al. 2014; Manser et al. 2015). However, there are also examples where mating preferences haven’t evolved against individuals carrying meiotic drive elements and theory suggests that in some cases this may not be expected (Price et al. 2012; Manser et al. 2017). The genetic system in *B. coprophila* is a little different to most meiotic drive systems, however. *Bradysia coprophila* doesn’t have one driving chromosome or allele (like for stalk eyed flies; Johns et al. 2005), but the entire maternal half of the genome in males exhibits drive, and the monogenic system may or may not be related to the drive system from an evolutionary standpoint. Therefore, it may be more difficult for mating preferences to evolve in this system, as the genetic basis of paternal genome elimination and monogeny may not be linked (Lande and Wilkinson, 1999). We are learning more about how monogeny is genetically encoded (Baird et al. 2023), however, there is still a limited understanding of the genetic basis of paternal genome elimination in this species, or any other species with this genetic system.

In some ways, the reproductive system in *B. coprophila* resembles the dynamics in hybridogenetic and gynogenetic species, where some individuals are of hybrid origin, and although they must mate to reproduce, they do not pass their partners genes onto their offspring. In these cases, like in *B. coprophila*, we expect individuals to avoid mating with gynogenetic or hybridogenetic individuals, as they will not pass their genes onto future generations. In hybridogenetic water frogs, for instance, like in our system, there is little evidence for male mate choice (Engeler and Reyer, 1998). However, in this system the lack of male mate preferences is interpreted as a male mate preference for larger females, as hybridogenetic females are often larger than conspecific females. In this species, although males show weak mating preferences against hybridogenetic partners, females do show strong mating preferences against hybridogenetic males (Engeler and Reyer, 1998). Mating preferences in males are expected to be harder to evolve than in females, as males have a steeper Bateman gradient (i.e. for males, mating with more mates increases their reproductive output to a greater extent than for females) (Arnold and Duvall, 1994). This may be part of the reason that we do not see mating preferences for gynogenic females in *B. coprophila*.

Furthermore, it has been suggested that the fact that males meet mates sequentially in the wild make it difficult for mating preferences to arise in males, and therefore, that studies should examine why males should refuse a mating opportunity rather than if he chooses to mate with one female over another (Barry and Kokko, 2010).

Mating dynamics in *B. coprophila* are also important for understanding population dynamics in this species. Male mate preferences could heavily influence the population sex ratio, as male fitness is entirely tied to the sex of his progeny. The lack of strong mate preferences in males makes sense from a population persistence perspective. If the mating preferences were strong in this species, we would expect populations to be unstable over time due to extreme sex ratio biases, which could lead to local population extinction. All Sciaridae species exhibit PGE as their reproduction system, but only some Sciaridae species are monogenic (Gerbi, 2022). It would be interesting to compare mating dynamics and ecology of monogenic vs. digenic species to understand whether these factors affect the evolution of mating preferences. For instance, we might expect monogenic systems to be stable only in species where ecological/ behavioural factors are such that male mating preferences for gynogenic vs. androgenic females is weak. For this reason, it might be interesting to study species with a mix of monogenic and digenic females, such as *B. ocellaris* (Metz 1938), as conflict between males and females over reproductive decisions might be higher than in monogenic species, where conflict may have already been resolved in favour of females controlling mating dynamics. We know very little about mating dynamics in Sciaridae species. We also have limited information about evolutionary transitions between monogeny and digeny in this clade, e.g. which of these systems is ancestral and if transitions can go either way.

Understanding these factors better would help answer questions about the evolution of monogeny and mating dynamics in Sciaridae. In addition to Sciaridae species, similar evolutionary dynamics also occur in gall gnats in the family Cecidomyiidae. Species in this family also exhibit PGE, and like Sciaridae, Cecidomyiidae also has a mix of monogenic species, digenic species, and species that have both monogenic and digenic females (White, 1973). Therefore, we would also expect males in monogenic Cecidomyiid species to be under selection to exhibit mate preferences towards gynogenic females. It would be interesting to see how mating preferences in Cecidomyiidae compares to what we see in Bradysia species.

Studies in these two clades provide a unique opportunity to study the interplay between the evolution of sexual conflict, sex determining mechanisms, and mate choice.

### Concluding remarks

*Bradysia coprophila* exhibits an intriguing system of reproduction that we would expect to generate male mating preferences, as males only gain fitness through mating with half the females in the population. We find however, that males exhibit very weak mating preferences, and that males seem to behave maladaptively and mate with androgenic females more often than gynogenic females. We propose a few explanations for these findings, namely that the mating dynamics we observed may be driven primarily by female receptivity rather than male preferences, as we expect the genetic system in this species to also influence how females will behave in mating interactions. Secondly, we suggest that securing mating partners for males may be more important than developing mate preferences against androgenic females. In any case, the stark contrast between the expectation of male mate preferences and the observed lack of preference is fascinating. Unraveling the factors that underlie the mating dynamics in this system in the future should provide insights into sexual selection and the evolution of preference in systems with meiotic drive.

## Acknowledgements

We would like to thank the Ross lab for useful suggestions throughout the duration of this experiment. We would also like to thank Gina Sinclair for help with preliminary experiments leading to this manuscript and Kamil Jaron, for comments on the manuscript.

## Data and resource availability

All data used in analyses are available on Zenodo (upon publication).

