## Supplementary material for "Does non-Mendelian chromosome transmission and unusual sex determination affect male mate choice in the fly *Bradysia coprophila*?"

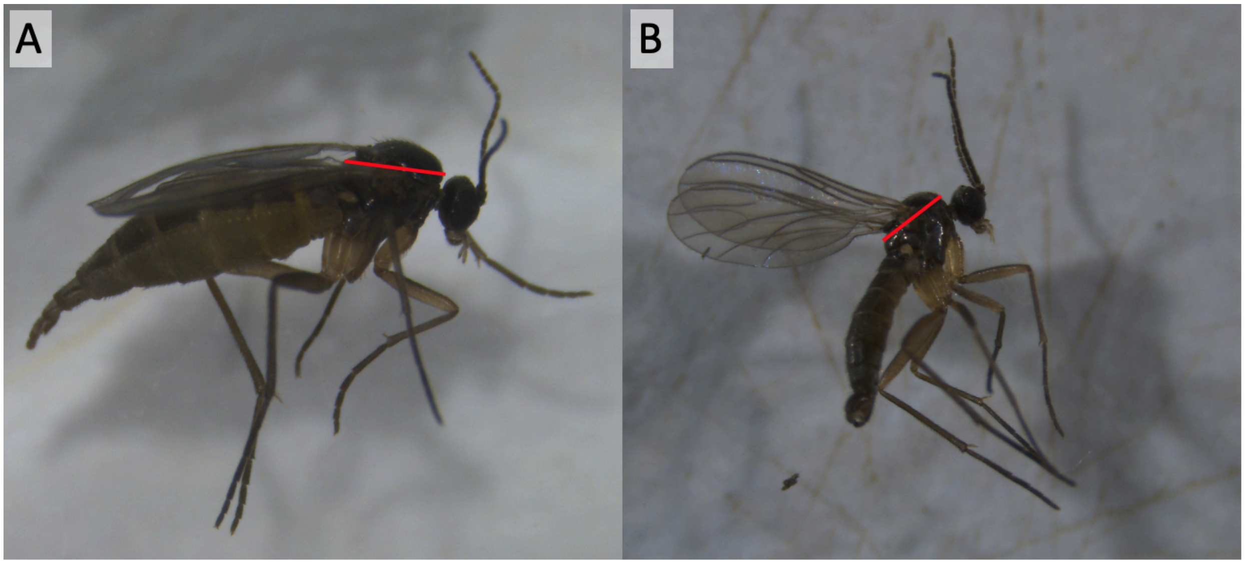

Supplementary Figure 1. Image of a *B. coprophila* female (A) and male (B) showing how we measured the thorax width (red line).

Supplementary Table 1. Sperm counts for males in the mating capacity experiment after mating. Males tended to have very few or many sperm left in their testes after mating.

| Fly line | Trial | Block | Density | Number of matings | Sperm count < 1000 | Sperm count <200 |
|---|---|---|---|---|---|---|
| Lab | T28 | 1 | Low | 6 | 85 | 85 |
| Lab | T32 | 1 | High | 6 | 6 | 6 |
| Lab | T34 | 1 | Low | 1 | 741 | 200 |
| Lab | T37 | 1 | High | 3 | 1000 | 200 |
| Lab | T39 | 1 | Low | 5 | 37 | 37 |
| Lab | T40 | 1 | Low | 5 | 139 | 139 |
| Lab | T43 | 1 | High | 7 | 527 | 200 |
| Lab | T45 | 1 | Low | 6 | 336 | 200 |
| Lab | T46 | 1 | High | 6 | 12 | 12 |
| Lab | T49 | 1 | Low | 7 | 1000 | 200 |
| Lab | T50 | 1 | High | 5 | 28 | 28 |
| Lab | T1 | 2 | High | 5 | | 0 |
| Lab | T2 | 2 | Low | 8 | | 0 |
| Lab | T5 | 2 | Low | 7 | | 39 |
| Lab | T7 | 2 | High | 6 | | 200 |
| Lab | T8 | 2 | Low | 4 | | 200 |
| Lab | T9 | 2 | High | 5 | | 200 |
| Lab | T10 | 2 | Low | 4 | | 200 |
| Lab | T11 | 2 | Low | 6 | | 50 |
| Lab | T12 | 2 | High | 6 | | 13 |
| Lab | T13 | 2 | High | 1 | | 200 |
| Lab | T14 | 2 | High | 6 | | 50 |
| Wild | CH12 | 1 | High | 3 | 1000 | 200 |
| Wild | CH15 | 1 | High | 8 | 165 | 165 |

| | | | | | | |
|---|---|---|---|---|---|---|
| Wild | CH20 | 1 | High | 5 | 25 | 25 |
| Wild | CH21 | 1 | High | 5 | 1000 | 200 |
| Wild | CH5 | 1 | High | 3 | 23 | 23 |
| Wild | CH6 | 1 | High | 4 | 1000 | 200 |
| Wild | CH7 | 1 | High | 5 | 100 | 100 |

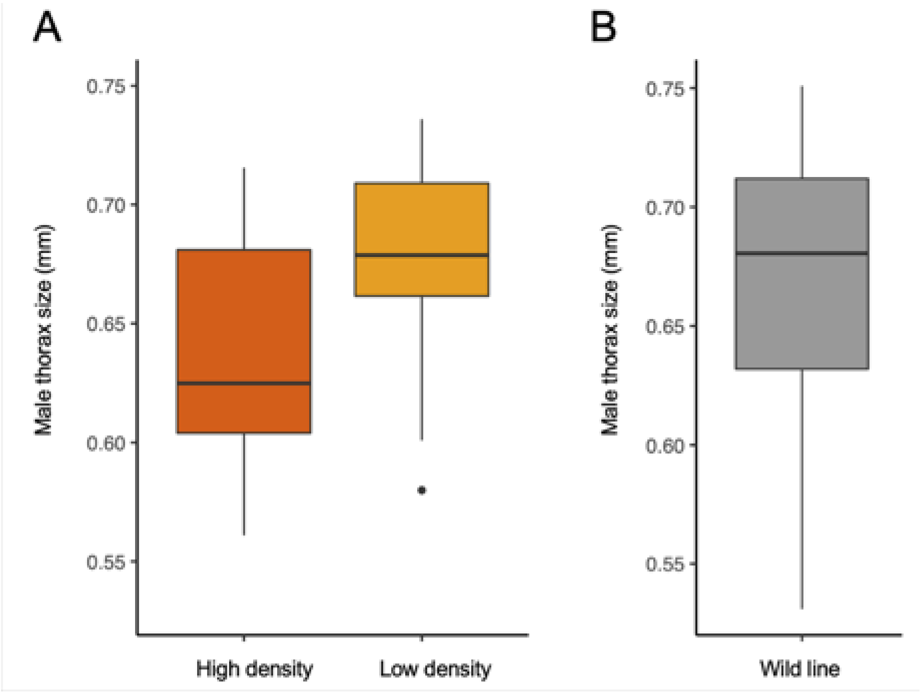

Supplementary Figure 2. (A) Thorax size differences between males from the lab line in high density (dark orange) and low density (light orange) treatments.(B) Size of males from the wild fly line.

Supplementary Table 2. Model estimates and standard error (in parentheses) for binary model analysing for flies from the lab line whether females have a different likelihood of mating based on their type (gynogenic/ androgenic) and male condition (low density/ high density upbringing). Male ID is added as a random effect in the model with the number of levels shown in parentheses and the variation and standard deviation (in parentheses) shown.

| | **Female mated (yes, no)** |
|---|---|
| **Fixed** | |
| Density | -0.121 (0.315) |
| Female Type | 0.558 (0.284) * |
| Interaction (Density*Female Type) | -0.466 (0.411) |
| Constant | -0.060 (0.217) |
| **Random** | |
| Male ID (42) | 0.173 (0.416) |
| Observations | 404 |
| Log Likelihood | -275.39 |
| AIC | 560.779 |
| BIC | 580.786 |
| Note: *p<0.05, **p<0.01, ***p<0.001 | |

Supplementary Table 3. Model estimates and standard error (in parentheses) for the model analysing for flies from the wild line whether males mated with different numbers of females of each type (gynogenic/ androgenic).

| | **Number of females mated** |
|---|---|
| **Fixed** | |
| Female type | -0.074 (0.193) |
| Constant | 0.981 (0.134) *** |
| Observations | 42 |
| Log Likelihood | -73.556 |
| AIC | 151.112 |
| Note: *p<0.05, **p<0.01, ***p<0.001 | |

Supplementary Table 4. Model estimates and standard error (in parentheses) for the poisson model analysing for flies from the lab line whether males produce a different number of male and female offspring (measured by whether the male produced more offspring with gynogenic or androgenic females) and whether male condition (low density/ high density upbringing) affects number of offspring of each sex. Male ID and observation number are added as random effects in the model with the number of levels shown in parentheses and the variation and standard deviation (in parentheses) shown.

| | **Number of larvae males produced** |
|---|---|
| **Fixed** | |
| Density | -0.634 (0.397) |
| Female Type | 0.336 (0.272) |
| Interaction (Density*Female Type) | -0.098 (0.397) |
| Constant | 4.458 (0.273) *** |
| **Random** | |
| Observation (84) | 0.786 (0.886) |
| Male ID (42) | 0.816 (0.904) |
| Observations | 84 |
| Log Likelihood | -498.651 |
| AIC | 1009.302 |
| BIC | 1023.886 |

Note: *p<0.05, **p<0.01, ***p<0.001

Supplementary Table 5. Model estimates and standard error (in parentheses) for the poisson model analysing for flies from the wild line whether males produce a different number of male and female offspring (measured by whether the male produced more offspring with gynogenic or androgenic females). Male ID and observation number are added as random effects in the model with the number of levels shown in parentheses and the variation and standard deviation (in parentheses) shown.

| | **Number of larvae males produced** |
|---|---|
| **Fixed** | |
| Female type | -0.262 (0.404) |
| Constant | 4.643 (0.294) *** |
| **Random** | |
| Observation (42) | 1.673 (1.294) |
| Male ID (21) | 0.119 (0.344) |
| Observations | 42 |
| Log Likelihood | -263.507 |
| AIC | 535.014 |
| BIC | 541.965 |

Note: *p<0.05, **p<0.01, ***p<0.001

Supplementary Table 6;. Model estimates and standard error (in parentheses) for the poisson model analysing for flies from the lab line whether the number of offspring a female produces depends on her type (gynogenic/ androgenic) and size (i.e. thorax width). Male ID and observation number are added as random effects in the model with the number of levels shown in parentheses and the variation and standard deviation (in parentheses) shown.

| | **Number of larvae females produced** |
|---|---|
| **Fixed** | |
| Female size | 22.167 (7.131) ** |
| Female type | 0.759 (0.633) |
| Constant | -16.060 (5.188) ** |
| **Random** | |
| Observation (183) | 10.99 (3.314) |
| Male ID (19) | 2.28 (1.510) |
| Observations | 183 |
| Log Likelihood | -622.738 |
| AIC | 1255.476 |
| BIC | 1271.523 |

Note: *p<0.05, **p<0.01, ***p<0.001

Supplementary Table 7. Model estimates and standard error (in parentheses) for the poisson model analysing for flies from the wild line whether the number of offspring a female produces depends on her type (gynogenic/ androgenic) and size (i.e. thorax width). Male ID and observation number are added as random effects in the model with the number of levels shown in parentheses and the variation and standard deviation (in parentheses) shown.

| | **Number of larvae females produced** |
|---|---|
| **Fixed** | |
| Female size | 2.426 (1.194) * |
| Female type | 0.159 (0.139) |
| Constant | 1.835 (0.952) |
| **Random** | |
| Observation (64) | 0.197 (0.444) |
| Male ID (15) | 0.338 (0.581) |
| Observations | 64 |
| Log Likelihood | -303.435 |
| AIC | 616.87 |
| BIC | 627.665 |

Note: *p<0.05, **p<0.01, ***p<0.001

Supplementary Table 8. Model output for poisson model measuring whether the number of females a male mated with affected the amount of sperm the male has left after the mating trial in the male mating capacity experiment. Values for each factor are the estimate and standard error.

| | **Sperm in testes after mating** |
|---|---|
| **Fixed** | |
| Number of matings | -0.385 (0.188) * |
| Constant | 6.008 (1.025) *** |
| **Random** | |
| Observation | 2.23 (1.493) |
| Observations | 22 |
| Log Likelihood | -128.62 |
| AIC | 263.239 |
| BIC | 266.512 |

Note: *p<0.05, **p<0.01, ***p<0.001

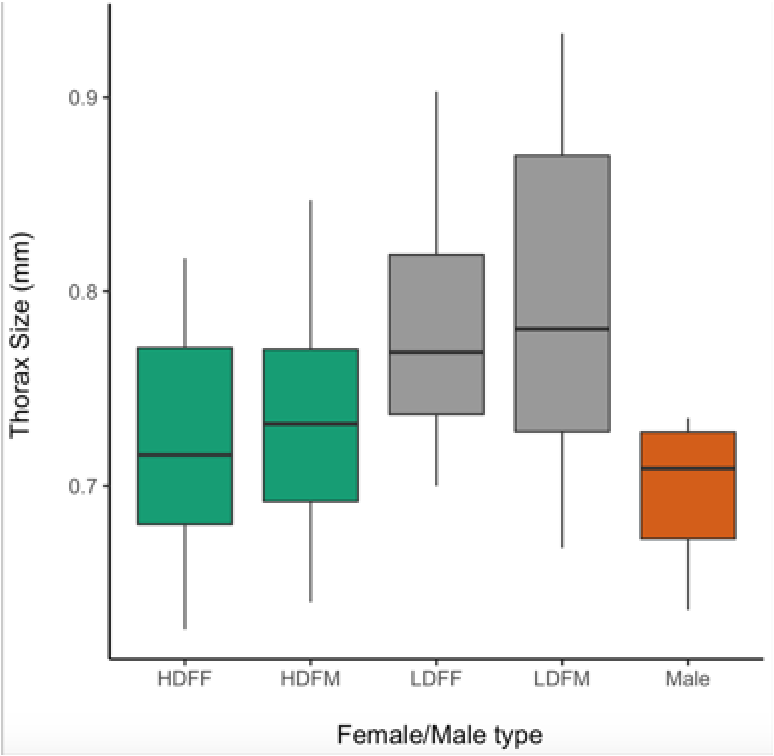

Supplementary Figure 3. Size differences between all female types (green or grey bars) and males (orange bar) in the mating preference experiment. The first two letters for the female categories indicate the density treatment (HD= high density (green bars), LD= low density (grey bars)), and the second two letters indicate the female type (FF= gynogenic or female producing, FM= androgenic or male producing).

Supplementary Table 9: Output statistics from the binary model analysing whether males in the mating preference experiment mated with females more based on their size (density) and type (gynogenic/ androgenic). Males mated with low density females and androgenic females more often. Values for each factor are the estimate and standard error.

| | **Mated female (yes/no)** |
|---|---|
| **Fixed** | |
| Density | 1.719 (0.817) * |
| Female Type | 1.917 (0.825) * |
| Interaction (Density*Female Type) | -1.254 (1.118) |
| Order of trial | 0.262 (0.259) |
| Constant | -1.501 (0.951) |
| **Random** | |
| Male ID (22) | 2.38 (1.54) |
| Observations | 88 |
| Log Likelihood | -51.305 |
| AIC | 114.61 |
| BIC | 129.474 |

Note: *p<0.05, **p<0.01, ***p<0.001

Supplementary Table 10: Output statistics from the binary model analysing whether females in the mating preference experiment resisted male mating based on their type and density treatment. Values for each factor are the estimate and standard error.

| | Female resisted mating (yes, no) |
|---|---|
| **Fixed** | |
| Density | 0.419 (0.533) |
| Female Type | 0.695 (0.540) |
| Constant | -0.0181 (0.576) |
| **Random** | |
| Male ID (22) | 2.523 (1.588) |
| observation (88) | 6.57e-07 (0.0008) |
| Observations | 88 |
| Log Likelihood | -54.505 |
| AIC | 119.01 |
| BIC | 131.396 |
| Note: *p<0.05, **p<0.01, ***p<0.001 | |

Supplementary Table 11: Poisson model output analysing whether males attempted to mate more (using number of thrusting attempts as a proxy) with females based on their size (density) and type (gynogenic/ androgenic).

| | **Number of thrusts towards female** |
|---|---|
| **Fixed** | |
| Female Type | 0.159 (0.362) |
| Density | 0.252 (0.361) |
| Observer | -0.373 (0.272) |
| Interaction (Density*Female Type) | -0.049 (0.505) |
| Constant | 1.72 (0.489) *** |
| **Random** | |
| Observation | 1.080 (1.039) |
| Male ID (22) | 0.054 (0.232) |
| Observations | 88 |
| Log Likelihood | -259.092 |
| AIC | 532.184 |
| BIC | 549.525 |

Note: *p<0.05, **p<0.01, ***p<0.001

Supplementary Table 12. Model output from the mixed effects model analysing whether the mating duration was affected by female type (gynogenic/ androgenic) or size (density treatment). The order in which males were presented with their partner was the most important factor in determining mating duration.

| | Mating Duration |
|---|---|
| **Fixed** | |
| Density | 0.070 (0.041) |
| Female Type | 0.020 (0.043) |
| Interaction (Density*Female Type) | -0.119 (0.056) * |
| Order of trial | 0.074 (0.012) *** |
| Constant | 2.638 (0.058) *** |
| **Random** | |
| Male ID (20) | 0.029 (0.170) |
| Residual | 0.008 (0.087) |
| Observations | 53 |

Note: *p<0.05, **p<0.01, ***p<0.001

Supplementary Table 13 : Poisson model output analysing whether the amount of sperm transferred to each female depended on female type (gynogenic/ androgenic) or size (density treatment).

| | **Number of sperm in spermatheca** |
|---|---|
| **Fixed** | |
| Female Type | 0.190 (0.194) |
| Density | 0.326 (0.179) |
| Order | 0.123 (0.078) |
| Mating duration | 0.001 (0.0004) |
| Interaction (Density*Female Type) | -0.392 (0.234) |
| Constant | 5.607 (0.339) *** |
| **Random** | |
| Observation | 0.050 (0.223) |
| Male ID (22) | 0.418 (0.647) |
| Observations | 27 |
| Log Likelihood | -171.978 |
| AIC | 359.956 |
| BIC | 370.322 |

Note: *p<0.05, **p<0.01, ***p<0.001
